# Key determinants of VDAC-hexokinase I complex assembly revealed by a minimal vesicle-based interaction assay

**DOI:** 10.1101/2025.11.03.686449

**Authors:** Milena Wessing, Isabelle Watrinet, Michael Timme, Britta Fiedler, Manel N. Melo, Jacob Piehler, Joost C. M. Holthuis

## Abstract

Binding of hexokinase HKI to mitochondrial voltage-dependent anion channels (VDACs) regulates the metabolic fate of glucose and promotes cell survival in hyperglycolytic tumors. Computer simulations indicated that complex assembly relies on intimate contacts between the *N*-terminal α-helix of HKI and a charged bilayer-facing glutamate on the outer wall of VDACs. Protonation of this residue blocks complex formation in silico, explaining the release of HKI from mitochondria observed upon cytosolic acidification. To validate these findings, we here developed an *in vitro* assay for interrogating HKI binding to VDAC1 reconstituted in vesicles captured on a functionalized surface. TIRF-based quantitative interaction studies with a fluorescent peptide comprising the *N*-terminal α-helix of HKI revealed a crucial role of the bilayer-facing glutamate in complex assembly and recapitulated an exquisite sensitivity of HKI-VDAC binding to fluctuations in pH. Our assay opens up important opportunities to investigate the impact of membrane environment on HKI-VDAC complex assembly and may benefit the development of therapeutics that target pathogenic imbalances in this process.

## INTRODUCTION

Voltage-dependent anion channels (VDACs) are abundant β-barrel proteins in the outer mitochondrial membrane (OMM) with crucial roles in mitochondrial physiology. While primarily known as gatekeepers for the passage of ions and metabolites like ATP/ADP and NAD+/NADH across the OMM [1, 2], VDACs participate in numerous other mitochondrial regulatory processes such as apoptosis [3-5], pyroptosis [6], calcium homeostasis [7], and lipid scrambling [8]. In mammalian cells, three isoforms exist (VDAC 1-3), each of which is able to mediate transport of ions and metabolites, but otherwise serve non-redundant functions through interactions with different protein binding partners [9, 10].

VDAC1 and VDAC2 have been implicated as dynamic binding platforms for the pro-apoptotic Bcl-2 proteins Bax and Bak [5, 11, 12], which oligomerize into pores that perforate the OMM to release cytochrome *c* and initiate caspase-mediated cell death [13]. Mitochondrial function and apoptotic cell death are also tied to VDAC oligomerization, which is strongly modulated by alterations in the membrane lipid environment [14]. Moreover, we previously showed that ceramides, the central intermediates of sphingolipid metabolism, can exert their apoptogenic activities directly on mitochondria [15, 16]. A chemical screen for mitochondrial ceramide binding proteins combined with molecular dynamics simulations and functional studies in cancer cells identified VDAC2 as critical effector of ceramide-induced apoptosis [17]. Ceramide binding critically relies on a conserved bilayer-facing glutamate on the outer channel wall. Mutation of this glutamate in VDAC2 abolished ceramide binding in simulations and rendered cancer cells resistant to ceramide-induced apoptosis [17].

VDAC1 and VDAC2 also serve as receptors for hexokinases (HKs). These enzymes catalyze phosphorylation of glucose to yield glucose-6-phosphate, the initial and rate-limiting step in glucose metabolism [18, 19]. Elevated levels of the mitochondrially bound HK isoforms HKI and HKII stimulate the rate of glycolysis and lactate production, causing a metabolic switch referred to as the Warburg effect [20]. Besides controlling the metabolic fate of glucose [21], VDAC-bound HKs confer apoptosis resistance in cancer cells by diminishing the propensity of VDACs to interact with Bax and Bak [22-25]. HK binding to VDACs is thought to play a pivotal role in promoting cell growth and survival in hyperglycolytic tumors [26, 27]. A reduced HKI interaction with VDACs has been implicated as causal factor in demyelinating peripheral neuropathies [28, 29]. Consequently, the binary protein complex has been recognized as potential target for therapeutic interventions [30-32].

HKI and HKII each contain a short *N*-terminal α-helix comprising 20 amino acids that harbors the VDAC binding site [33, 34]. Previous modeling studies yielded conflicting models of how HKs and VDACs assemble into a binary complex [35-37] and failed to address the essential role of the bilayer-facing glutamate located on the outside wall of VDAC1 in HKI binding [38, 39]. Using molecular dynamics [MD] simulations and functional studies in cells, we recently identified the core structural and topological determinants that govern HKI binding to VDAC1 and VDAC2 [40]. We showed that complex assembly relies on intimate contacts between the *N*-terminal α-helix of HKI and the bilayer-facing glutamate of VDAC1 and VDAC2. A negatively charged, deprotonated glutamate turned out to be crucial for HKI binding. Hence, we observed that protonation of the bilayer-facing glutamate abolished HKI binding in simulations while transient acidification of the cytosol caused a reversable release of a *N*-terminal HKI peptide from mitochondria [40]. Our finding that anti-apoptotic HKI and pro-apoptotic ceramides share a common binding site on the outer wall of VDACs points at a potential mechanism by which ceramides exert their apoptogenic activities.

Here, we report the development of a minimal vesicle-based protein-protein interaction platform to reconstitute HKI-VDAC1 complex assembly *in vitro*. Co-reconstitution of VDAC1 with a maltose binding protein fused to a transmembrane domain allowed immobilization of VDAC1-containing vesicles to a glass substrate decorated with αMBP-DARPin. Subsequent TIRF-based quantitative binding studies with a fluorescent peptide comprising the *N*-terminal α-helix of HKI enabled experimental validation of a critical role of the bilayer-facing glutamate in complex assembly and recapitulated the exquisite sensitivity of VDAC-HKI binding to fluctuations in pH. Thus, we established a novel and versatile approach to investigate how the membrane lipid environment influences HKI-VDAC complex assembly and facilitate the development of therapeutic compounds aimed at modulating this process.

## RESULTS

### HKI-N binding to VDAC1 is controlled by protonation of a membrane-buried Glu

HKI contains an *N*-terminal α-helix of 17 residues (HKI-N) that is both essential and sufficient to bind mitochondrial VDACs [40]. For MD simulations of HKI-N binding to VDACs, we used an α-helical peptide comprising HKI-N with an additional Gln at its *C*-terminus (corresponding to Gln18 in HKI). Our initial simulations of HKI-N binding to VDACs were performed in an OMM-mimicking bilayer. The bilayer composition was based on Horvath and Daum [41] and featured an asymmetric transbilayer distribution of phosphatidylinositol and phosphatidylethanolamine [40]. As biochemical reconstitution of VDACs in an OMM-mimicking lipid environment would be technically very challenging, we first tested whether HKI-N binding to VDACs simulated in a simple, 1-palmitoyl-2-oleoyl-*sn*-glycero-3-phosphocholine (POPC) bilayer is governed by the same structural and physicochemical features as in our previous simulations. VDAC1 embedded in a POPC bilayer readily formed stable contacts with HKI-N, provided that the channel’s *C*-terminus faced the IMS-mimicking leaflet and the membrane-buried Glu73 (E73) was deprotonated (**Fig. 1a-d**). Under these conditions, the *N*-terminal half of HKI-N was able to insert vertically into the cytosolic membrane leaflet and slide along the channel wall to make direct contact with the negatively charged Glu73 (E73^−^). These contacts were primarily mediated by HKI-N residues Met1 and Gln5, both situated on the same side along the axis of the α-helix (**Fig. 1e**). Protonation of the membrane-buried Glu73 or substitution of Gln for Glu73 in each case abolished HKI-N binding (**Fig. 1d, e**). These results are in line with our previous simulations in OMM-mimicking bilayers, indicating that the protonation state of the membrane-facing Glu on the outer wall of VDAC1 is a critical determinant of HKI-N binding and that complex formation does not require an OMM-specific lipid environment but also occurs in a simple PC bilayer.

**Figure 1.**
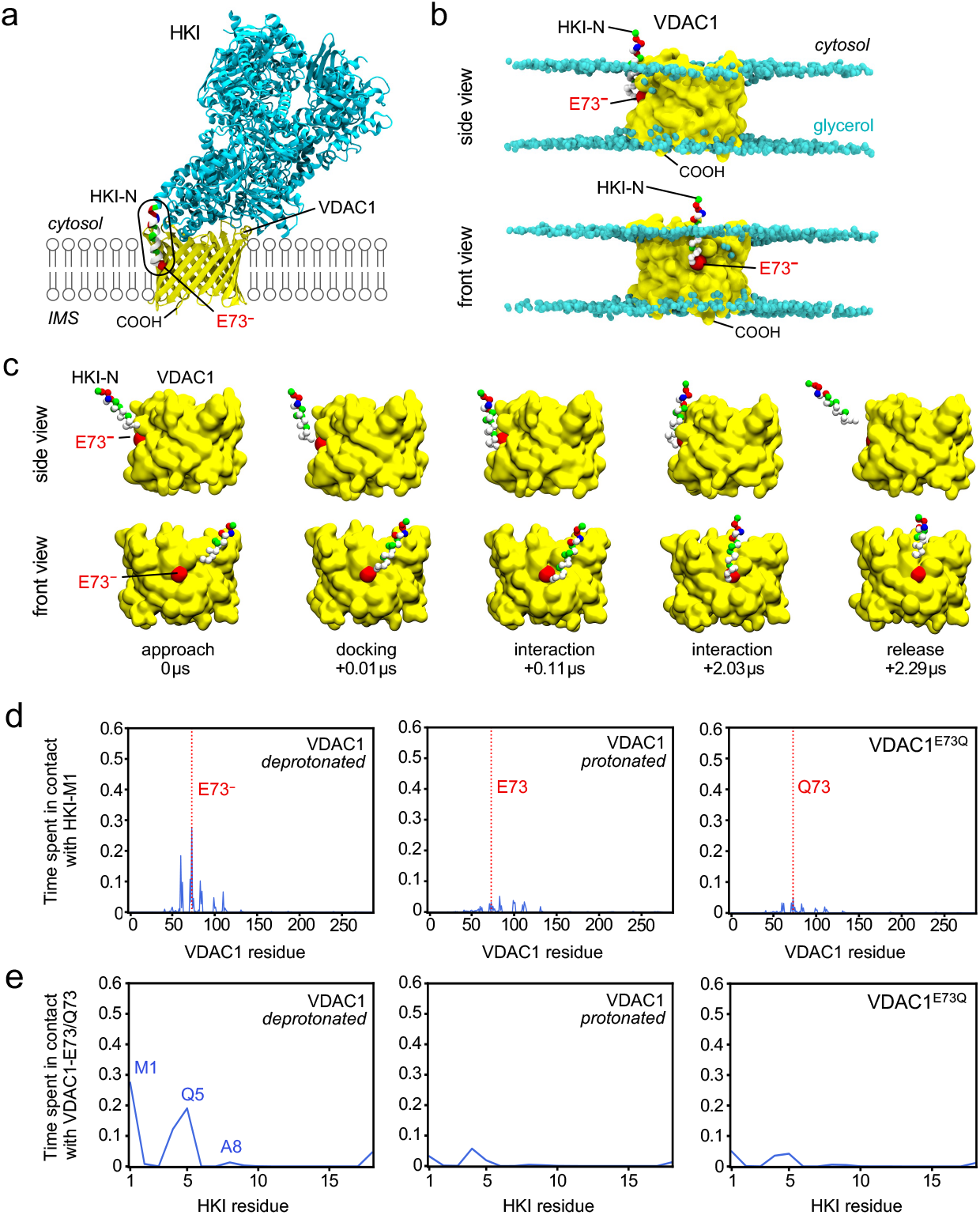
MD simulations of VDAC1–HKI-N complex assembly in PC bilayers. **(a)** Atomic model of HKI (*cyan*, with residue-type colored HKI-N) bound to VDAC1 (*yellow*) with the membrane-buried Glu (E73) marked in red. Image reproduced from [40]. **(b)** Stills from an MD simulation showing HKI-N bound to VDAC1 with a deprotonated E73 (*red*) and IMS-facing *C*-terminus. Glycerol groups in the POPC bilayer are marked in *cyan*. **(c)** Stills from an MD simulation, showing the approach and binding of HKI-N to VDAC1 with a deprotonated E73 (*red*) and IMS-facing *C*-terminus (away from the cytosol). **(d)** Relative duration of contacts between HKI-Met1 and specific residues of VDAC1 with a protonated or deprotonated E73 and VDAC1^E73Q^. All simulations were carried out in a POPC bilayer with the *C*-termini of VDAC1 facing the IMS. Shown are the combined data of three individual replicas with a total simulation time between 274 μs and 334 μs per condition. **(e)** Relative duration of contacts between VDAC1-E73 or VDAC1-Q73 and specific residues of HKI-N under the same conditions as in (d).

### Design of a minimal vesicle-based VDAC1–HKI-N interaction assay

HKI binding to mitochondrial VDACs promotes the glycolytic state of cancer cells while antagonizing activation and oligomerization of pro-apoptotic Bcl-2 proteins in response to apoptotic cues. This notion raised considerable interest in small anticancer compounds that perturb HKI-VDAC interactions. Development of such compounds requires assays in which these interactions can be faithfully reproduced. With this in mind, we designed a minimal surface-based assay in which the binding kinetics of a fluorescently-labeled HKI-N peptide to VDAC1 reconstituted into vesicles can be monitored by FRAP (**Fig. 2a**). Co-reconstitution of MBP with an artificial transmembrane domain (MBP-TMD; **Suppl. Fig. 1**) was employed for selective surface capturing via an immobilized αMBP-DARPin. To this end, we devised surface coating with a highly biocompatible PEG polymer brush that was site-specifically functionalized with the αMBP-DARPin via a *C*-terminal cysteine residue. To validate the functionality of the αMBP-DARPin-modified surface, the coverslips were incubated for 30 min at RT with recombinant mEGFP or mEGFP fused to MBP (MBP-mEGFP), washed, and analyzed by TIRF microscopy for mEGFP fluorescence. This revealed that MBP-mEGFP, but not mEGFP was selectively bound to the αMBP-DARPin-modified surface (**Fig. 2b, d**). Vesicle capturing to these surfaces was probed using MBP-TMD reconstituted at a protein:lipid ratio of 1:500 into very small unilamellar vesicles composed of 99.5 mol% L-α-phosphatidylcholine (egg PC) and 0.5 mol% Oregon Green 488-labeled 1,2-dihexadecanoyl-*sn*-glycero-3-phosphoethanolamine (OG488-DHPE) as fluorescent vesicle marker. We used methyl-β-cyclodextrin for detergent depletion to obtain homogeneous very small unilamellar vesicles, which can be directly used for surface capturing [42]. Binding experiments with protein-free or MBP-TMD-containing vesicles followed by TIRF microscopy analysis for OG488 fluorescence showed that only the latter where effectively captured on the αMBP-DARPin-modified surface (**Fig. 2c, d**), confirming that αMBP-DARPin coupled to the glass surface retained its functionality. In FRAP experiments with MBP-TMD containing vesicles bound to the functionalized surface, we could not detect any measurable recovery of OG488 fluorescence in the bleached area over a prolonged period of time (150 sec; **Fig. 2e**). This indicated that vesicles captured on the modified surface remain largely intact and do not undergo fusion to form a membrane bilayer. With this, an important requirement for meaningful vesicle-based VDAC1-HKI-N interaction studies was met.

**Figure 2.**
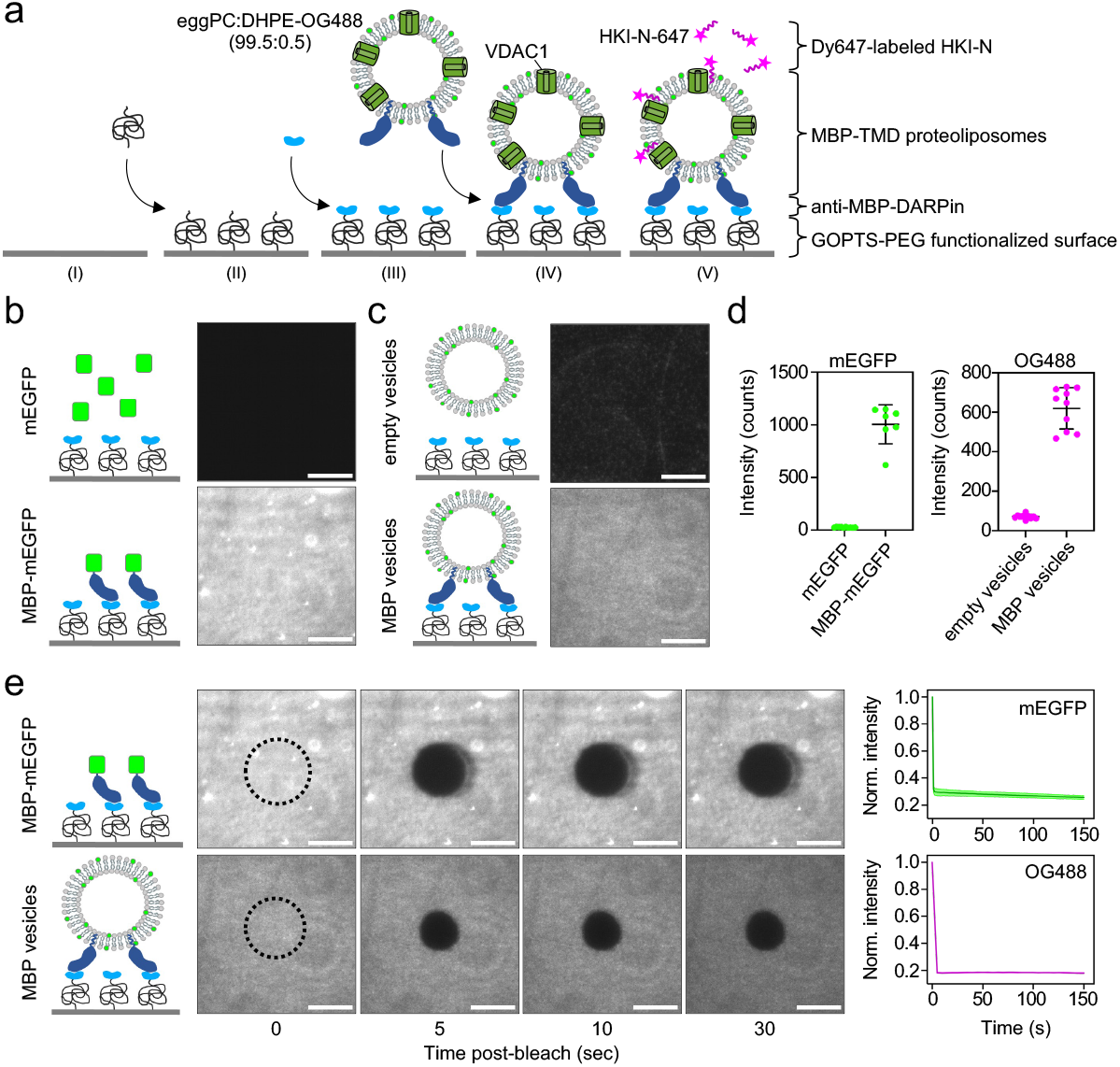
MBP-TMD-containing vesicles readily bind to αMBP-DARPin-modified surface and do not undergo membrane fusion. **(a)** Schematic outline of minimal vesicle-based protein interaction assay. Vesicles containing maltose binding protein fused to a transmembrane domain (MBP-TMD) are captured on glass coverslips functionalized with αMBP-DARPin via crosslinking to GOPTS-PEG-NHS maleimide. Co-reconstitution of VDAC1 gives rise to vesicles that bind DY-647P1-labeled peptide representing the *N*-terminal α-helix of HKI. OG488-labeled DHPE serves as fluorescent vesicle marker. **(b)** Coverslips functionalized with αMBP-DARPin as in (a) were incubated with mEGFP or MBP-mEGFP, washed, and analyzed for mEGFP fluorescence using TIRF microscopy. **(c)** Coverslips functionalized as in (b) were incubated with protein-free (empty) or MBP-TMD-containing vesicles, washed and analyzed for OG488 using TIRF microscopy. **(d)** Intensity plots of mEGFP and OG488 fluorescence at the surface of coverslips treated as in (b) and (c). **(e)** Photobleaching of MBP-mEGFP or MBP-TMD-containing vesicles captured on coverslips functionalized as in (b). Photobleaching was with a 405 nm laser for 30 sec followed by post-bleach imaging at 5 sec intervals over a period of 150 sec using a 488 nm laser. Images were acquired by TIRF microscopy. MBP-TMD was reconstituted at a molar protein:lipid ratio of 1:500 in eggPC:DHPE-OG488 (99.5:0.5). Data shown are representative of three independent experiments. Scale bar, 10 μm.

### VDAC1-containing vesicles are effectively captured to modified glass surface

We next included VDAC1 in MBP-TMD-containing vesicles and analyzed their capturing onto αMBP-DARPin-modified surfaces. To this end, single-cysteine variants of VDAC1 and VDAC1^E73Q^ were expressed in *E. coli*, purified by cation exchange chromatography, labeled with DY-547P1 maleimide, and subjected to size exclusion chromatography (**Suppl. Fig. 2**). VDAC channels are the most abundant proteins in the OMM, covering from 30% up to 80% of the membrane surface area in high-density regions [14, 43, 44]. This VDAC density corresponds to a molar protein:lipid ratio between 1:50 and 1:100. To obtain vesicles with a VDAC1 content similar to that of native OMM, we reconstituted unlabeled VDAC1 at a protein:lipid ratio ranging from 1:10 to 1:100 while keeping the protein:lipid ratio for MBP-TMD constant at 1:500. The efficiency by which these vesicles were captured on αMBP-DARPin-modified surfaces was assessed by TIRF microscopy using OG488 fluorescence as vesicle marker. Vesicles lacking VDAC1 and containing only MBP-TMD (empty vesicles) served as control. VDAC1-containing vesicles prepared with a protein:lipid ratio of 1:70 bound the functionalized surface with similar efficiency as empty vesicles. However, VDAC1-containing vesicles prepared with a protein:lipid ratio of 1:50 displayed a significant drop in surface binding and binding became also more uneven, presumably due to vesicle clustering (**Suppl. Fig. 3a, c**). For VDAC1^E73Q^-containing vesicles, binding to the surface was already significantly diminished when using a protein:lipid reconstitution ratio of 1:100 (**Suppl. Fig. 3b, c**). Immunoblot analysis revealed that equal amounts of vesicle-associated VDAC1 and VDAC1^E73Q^ could be captured on the surface when vesicles were prepared using protein:lipid reconstitution ratios of 1:50 for VDAC1 and 1:100 for VDAC1^E73Q^ (**Fig. 3b**). We also noticed that incorporation of VDAC1 reduced the amount of MBP-TMD present in the vesicles, which may explain why VDAC1-containing vesicles bind the functionalized surface less efficiently than empty vesicles. In line with the immunoblot data, TIRF microscopy of surface-bound vesicles prepared with DY-547P1-labeled VDAC1 and VDAC1^E73Q^ reconstituted at 1:50 and 1:100 protein:lipid ratios, respectively, revealed equal levels of DY-547P1 fluorescence at the surface (**Fig. 3c, d, f**). Therefore, we kept these ratios the same for the remainder of the study. In FRAP experiments, we observed no measurable recovery of either DY-547P1 (**Fig. 3d, e**) or OG488 fluorescence (**Fig. 3g-i**) in the bleached area over a prolonged period of time (300 sec). This indicates that the VDAC1- and VDAC1^E73Q^-containing vesicles remain tightly bound to the surface and do not undergo fusion to form a membrane bilayer.

**Figure 3.**
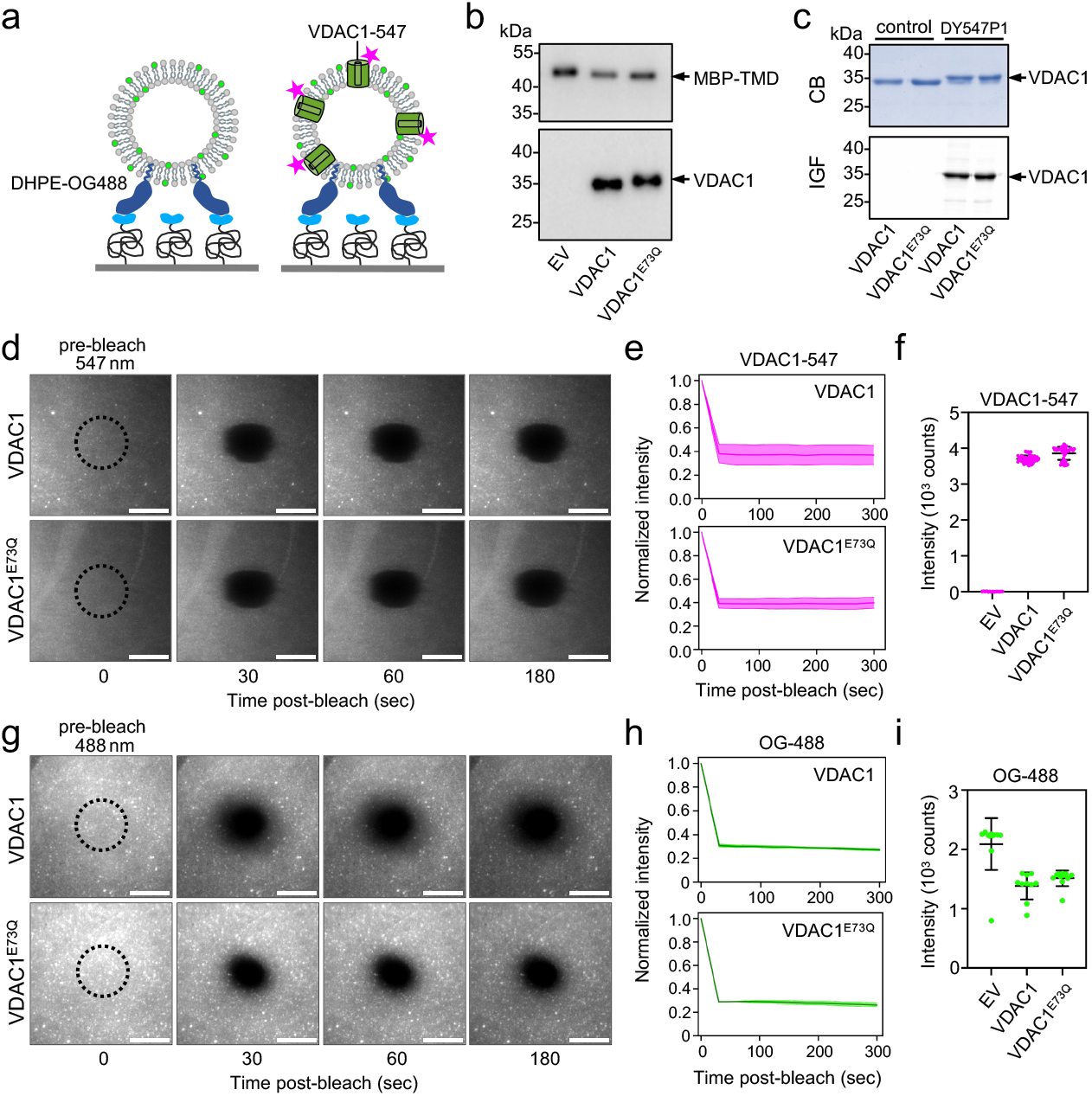
VDAC-containing vesicles immobilized on αMBP-DARPin-modified surface remain intact. **(a)** Cartoon of vesicles generated by (co)-reconstitution of MBP-TMD and DY-547P1-labeled VDAC1 and captured on coverslips functionalized with αMBP-DARPin. **(b)** Vesicles containing MBP-TMD in combination with VDAC1 or VDAC1^E73Q^ were extracted from αMBP-DARPin-functionalized coverslips and subjected to immunoblot analysis using antibodies against MBP and VDAC1. **(c)** Control and DY-547P1-labeled VDAC1 and VDAC1^E73Q^ used in reconstitution experiments were analyzed by SDS-PAGE, Coomassie Blue (CB) staining and in-gel fluorescence (IGF). **(d)** Vesicles containing MBP-TM and DY-547P1-labeled VDAC1 or VDAC1^E73Q^ were prepared as in (b), captured on αMBP-DARPin-functionalized coverslips and a circular ROI as indicated by the dotted lines was photobleached at 405 nm for 0.2 s. Post-bleach images were taken by TIRF microscopy at 30 s intervals over a period of 300 s using a 561 nm laser. Scale bar, 10 μm. **(e)** Recovery curves of photo-bleached vesicles as in (d). Data were corrected for imaging-related bleaching and plotted as normalized intensity over time. **(f)** Intensity plots of DY-547P1 fluorescence at the surface of coverslips treated as in (d). **(g)** Vesicles prepared as in (d) were captured on αMBP-DARPin-decorated coverslips and subjected to photobleaching for 0.2 s using a cellFRAP unit. Post-bleach images were taken by TIRF microscopy at 30 s intervals over a period of 300 s using a 488 nm laser. Scale bar, 10µm. **(h)** Recovery curves of photo-bleached vesicles as in (g). Data were corrected for imaging-related bleaching and plotted as normalized intensity over time. **(i)** Intensity plots of DHPE-OG488 fluorescence at the surface of coverslips treated as in (g). Vesicles containing MBP-TMD, VDAC1 and/or VDAC1^E73Q^ were prepared using a molar protein:lipid reconstitution ratio of 1:500 (MBP-TMD), 1:50 (VDAC1) and 1:100 (VDAC1^E73Q^) in eggPC:DHPE-OG488 (99.5:0.5). Data shown are representative of three independent experiments and plotted as mean values ± SD of 5-10 FRAPs per surface per condition.

### Binding of HKI-N to VDAC1 is disrupted by mutation of the bilayer-facing Glu73

MD simulations indicated that HKI-VDAC1 complex assembly in the OMM critically relies on intimate contacts between the *N*-terminal α-helix of HKI and a deprotonated, negatively charged membrane-buried Glu, E73, on the outer channel wall. To experimentally validate this finding under biochemically defined conditions, we quantified binding of a peptide corresponding to the first 25 amino acids of human HKI (HKI-N) to reconstituted VDAC1. To this end, surface-captured vesicles containing VDAC1, VDAC1^E73Q^ or only MBP-TMD, respectively, were incubated with HKI-N site-specifically labelled with DY-647P1 maleimide at Cys19 (HKI-N^647^). Peptide binding to the surface-captured vesicles was probed using TIRF microscopy and FRAP (**Fig. 4a**). After measurements, the MBP-TMD, VDAC1 and VDAC1^E73Q^ content of the vesicles was assessed by immunoblot analysis following their extraction from the surface (**Fig. 4b**). This revealed that HKI-N^647^ readily bound to VDAC1-containing vesicles (**Fig. 4c, e**). In contrast, peptide binding to VDAC1-deficient (empty) or VDAC1^E73Q^-containing vesicles was drastically reduced (**Fig. 4c, e**). Considering the fluorescence level associated with empty vesicles as background from the bulk solution, a 95% decrease of the FRAP on HKI-N^647^ bound to VDAC1-containing vesicles showed full fluorescence recovery, with a two-phasic characteristic (**Fig. 4c, f**). The initial, essentially instantaneous partial fluorescence recovery can be ascribed to bulk fluorescence, while the slower components can be related to the exchange kinetics of VDAC1-bound HKI-N^647^. In line with this interpretation, FRAP on HKI-N^647^ bound to empty vesicles revealed only the instantaneous fluorescence recovery background component. Strikingly, similarly fast fluorescence recovery was observed for HKI-N^647^ exposed to VDAC1^E73Q^, confirming much weaker binding as compared to VDAC1. Fitting of the slower component observed for the fluorescence recovery of HKI-N^647^ bound to VDAC1, a complex lifetime of 5-6 s was estimated. Consistent with our MD simulations (**Fig. 1c-e**), formation HKI-N binding was abolished in the absence of VDAC1 or by replacing the membrane-buried Glu (E73) on the outer channel wall for Gln. Collectively, these results indicate that our minimal vesicle-based protein binding assay is suitable for quantifying specific interactions between VDAC1 and HKI-N under biochemically defined conditions.

**Figure 4.**
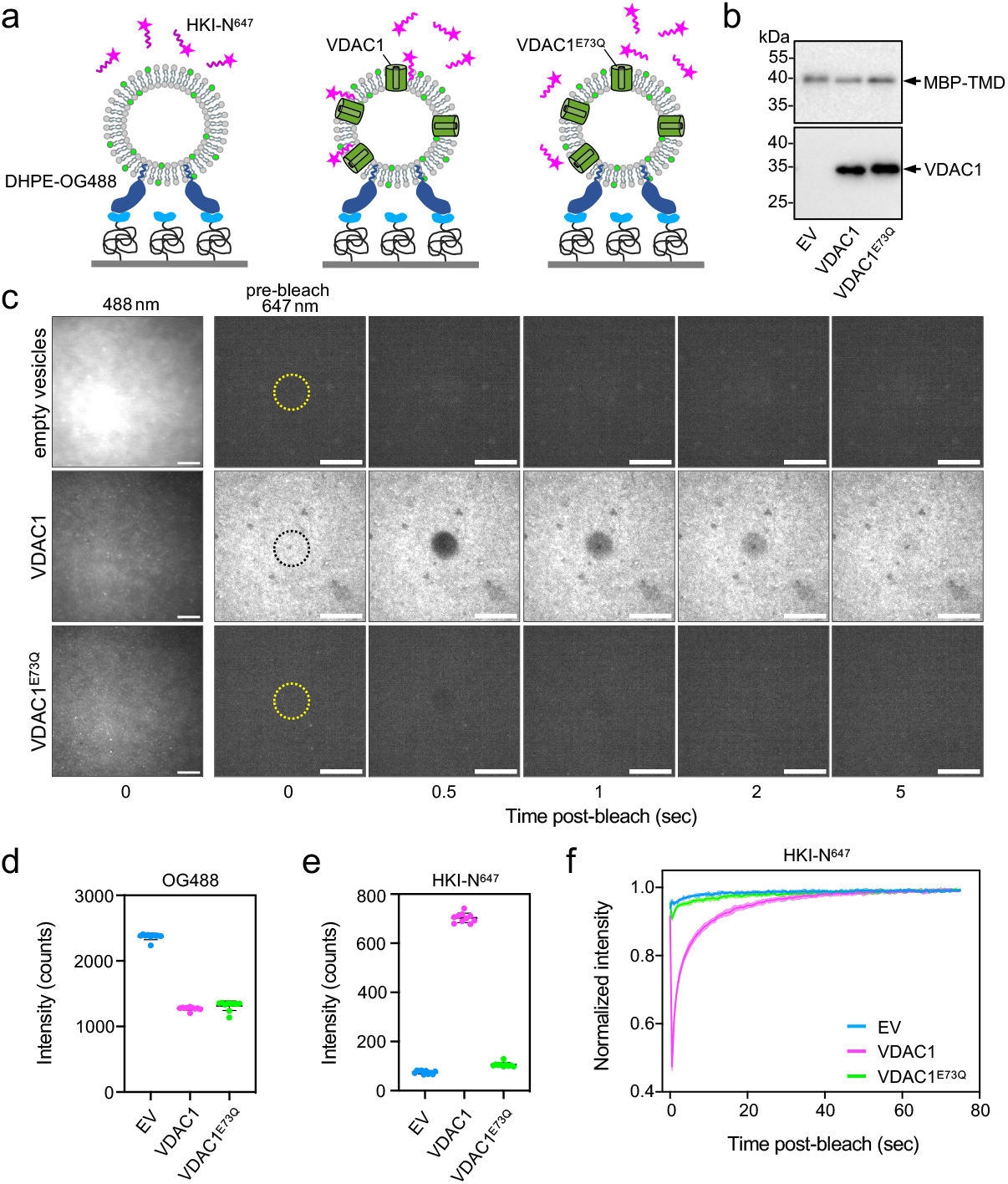
Substitution of Gln for Glu73 in VDAC1 abolishes HKI-N binding. **(a)** Schematic outline of VDAC1-HKI-N binding assay. Vesicles generated by (co)-reconstitution of MBP-TM, VDAC1 and/or VDAC1E73Q are captured on αMBP-DARPin-decorated coverslips and then analyzed for their ability to bind DY-647P1-labeled HKI-N peptide (HKI-N^647^) using TIRF microscopy. **(b)** Vesicles containing MBP-TMD in combination with VDAC1 or VDAC1^E73Q^ were extracted from αMBP-DARPin-decorated coverslips and subjected to immunoblot analysis using antibodies against MBP and VDAC1. **(c)** Vesicles bound to αMBP-DARPin-decorated coverslips as in (b) were incubated with 400 nM HKI-N^647^ for 5 min, and then a circular ROI indicated by the yellow dotted lines was photobleached at 405 nm for 0.2 s. Post-bleach fluorescence images were recorded at 0.5 s time intervals over a period of 75 s using TIRF excitation at 642 nm. Scale bar, 10 µm. **(d)** Intensity plots of DHPE-OG488 fluorescence at the surface of coverslips treated as in (c). **(e)** Intensity plots of HKI-N^647^ fluorescence at the surface of coverslips treated as in (c). Each dot represents a measurement **(f)** Recovery curves of HK-N^647^ fluorescence levels in areas of photo-bleached vesicles as in (c). Data were corrected for imaging-related bleaching and plotted as normalized intensity over time. Vesicles containing MBP-TMD, VDAC1 and/or VDAC1^E73Q^ were prepared using a molar protein:lipid reconstitution ratio of 1:500 (MBP-TMD), 1:50 (VDAC1) and 1:100 (VDAC1^E73Q^) in eggPC:DHPE-OG488 (99.5:0.5). Data shown are representative of three independent experiments and plotted as mean values ± SD of 5-10 FRAPs per surface per condition.

### Mild acidification causes an acute but reversible dissociation of the VDAC1–HKI-N complex

Protonation of Glu73 disrupts HKI-N binding in simulations while transient acidification of the cytosol causes a reversible release of HKI-N from mitochondria [40]. We therefore employed our minimal vesicle-based interaction assay to probe the impact of fluctuations in pH on HKI-N binding to VDAC1. To this end, vesicles containing VDAC1, VDAC1^E73Q^ or empty vesicles captured on a functionalized surface were incubated with HKI-N^647^ at pH 7.5 and analyzed for peptide binding using TIRF microscopy. As expected, only VDAC1-containing vesicles displayed robust peptide binding (**Fig. 5a, b**). Mild acidification of the incubation buffer from pH 7.5 to pH 6.5 drastically reduced the HKI-N^647^-fluorescence associated with the VDAC1-containing vesicles. This was not caused by an acidification-induced quenching of the fluorophore attached to the peptide since the bulk fluorescence levels associated with empty vesicles or vesicles containing VDAC1^E73Q^ remained unaltered upon lowering the pH. Mild acidification also had no impact on vesicle-associated OG488 fluorescence levels (**Suppl. Fig. 4**), ruling out that mild acidification somehow induced a release of vesicles from the functionalized surface. Upon raising the pH of the incubation buffer back to 7.5, binding of HKI-N^647^ to VDAC1-containing vesicles was largely restored (**Fig. 5a, b**). FRAP experiments revealed that at pH 6.5, the peptide fluorescence in the bleached area did not recover to the same degree as that observed at pH 7.5 (**Fig. 5c, d**). This could be due to the possibility that mild acidification results in protonation of E73 in many but not all VDAC1 channels. In any case, raising the pH back to 7.5 restored the characteristics of fluorescence recovery curves, confirming full reversibility of the effect as expected for protonation (**Fig. 5c, d**). Fitting of the fluorescence recovery curves revealed that mild acidification from pH 7.5 to pH 6.5 reduced the complex lifetime from ∼6s to ∼4s. The complex lifetime was restored to the original value when the pH was returned to 7.5 (**Fig. 5e**). FRAP on HKI-N^647^ bound to empty vesicles or vesicles containing VDAC1^E73Q^ did not yield differences in fluorescence recovery kinetics upon mild acidification (**Fig. 5d**), in line with the interpretation that fluorescence signals in these experiments are dominated by bulk fluorescence. Together, these results pinpoint pH-induced changes in VDAC1-HKI-N complex assembly previously observed in living cells [40] to protonation/deprotonation rather than potential indirect effects.

**Figure 5.**
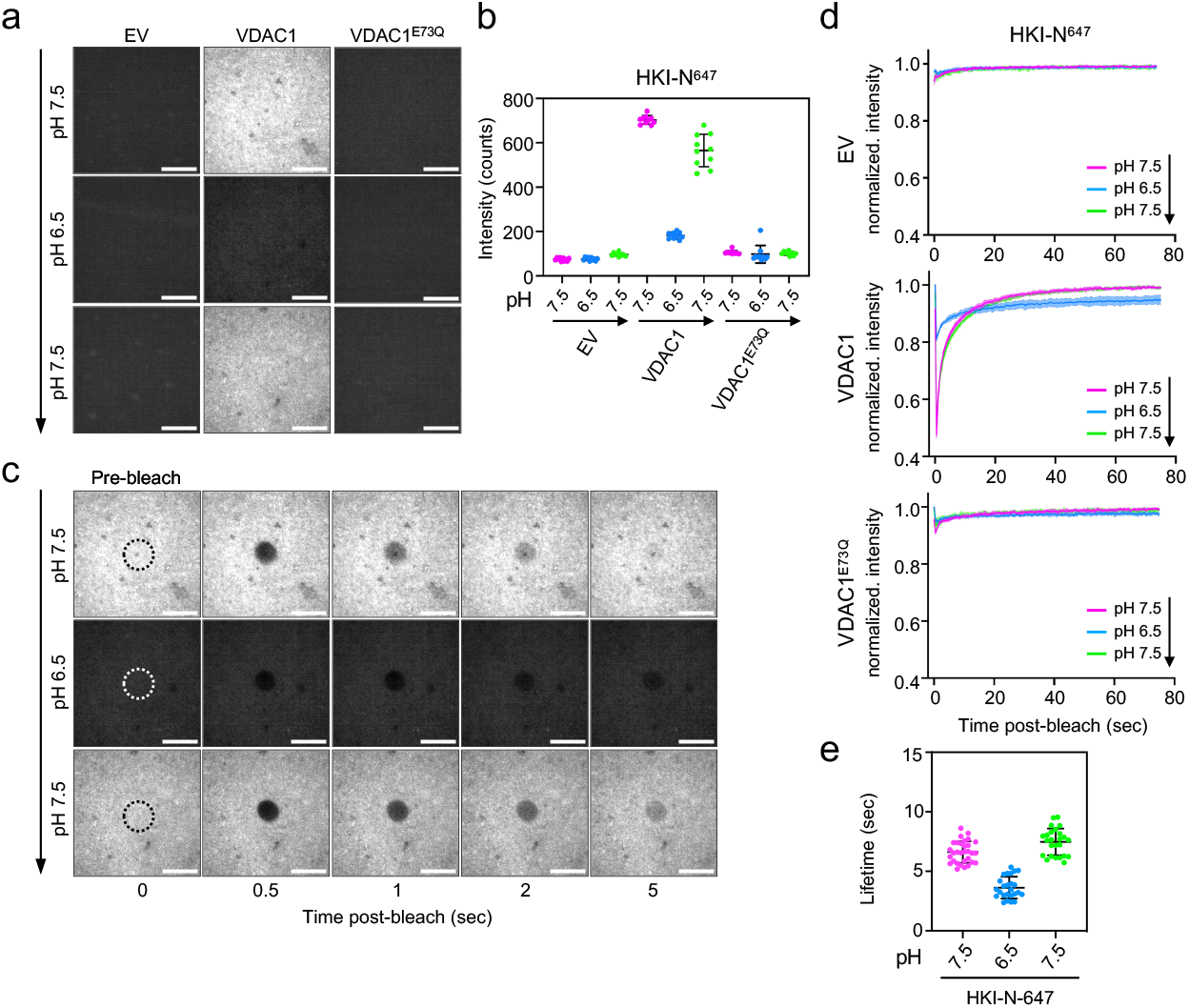
HKI-N binding to VDAC1 is pH sensitive. **(a)** Vesicles generated by (co)-reconstitution of MBP-TMD, VDAC1 and/or VDAC1E73Q were immobilized on αMBP-DARPin-decorated coverslips. Next, vesicles were incubated with 400 nM HKI-N^647^ for 5 min in pH 7.5 buffer and analyzed for peptide binding using TIRF microscopy. Incubation with HKI-N^647^was then continued in pH 6.5 buffer for 5 min before measuring peptide binding. This procedure was then repeated in pH 7.5 buffer. **(b)** Intensity plots of DY-647P1 fluorescence at the surface of coverslips treated as in (a). **(c)** Vesicles as in (a) were incubated with 400 nM HKI-N^647^ in buffer at the indicated pH and then a circular ROI indicated by the yellow dotted lines was photobleached at 405 nm for 0.2 s. Post-bleach fluorescence images were recorded at 0.5 s time intervals over a period of 75 s using TIRF excitation at 642 nm. Scale bar, 10µm. **(d)** Recovery curves of HK-N^647^ fluorescence in areas of photo-bleached vesicles as in (c). Data were corrected for imaging-related bleaching and plotted as normalized intensity over time. **(e)** Lifetimes of HK-N^647^-VDAC1 interactions measured in buffers at the indicated pH. The lifetimes were determined using a mono exponential decay over the first 25 sec (for pH 6.5 the first 10 sec) after photobleaching. Vesicles containing MBP-TMD, VDAC1 and/or VDAC1^E73Q^ were prepared using a molar protein:lipid reconstitution ratio of 1:500 (MBP-TMD), 1:50 (VDAC1) and 1:100 (VDAC1^E73Q^) in eggPC:DHPE-OG488 (99.5:0.5). Data shown are representative of three independent experiments and plotted as mean values ± SD of 5-10 FRAPs per surface per condition.

## DISCUSSION

HKI binding to mitochondrial VDACs plays a crucial role in controlling the metabolic fate of glucose and shifts the susceptibility of mitochondria to pro-apoptotic signals mediated through Bcl2-family members. Perturbation of HKI-VDAC complex assembly has been recognized as potential therapeutic strategy to curb cell growth and survival in rapidly growing tumors. We recently uncovered the structural basis of HK-VDAC complex formation using MD simulations and provided evidence that HKI directly binds to a charged bilayer-facing Glu on the outer channel wall via its *N*-terminal amphipathic α-helix [40]. We here developed a surface-based in *vitro* assay to directly interrogate HKI binding to VDAC1 reconstituted into a well-defined membrane context. We observed that the interaction of a peptide comprising the *N*-terminal α-helix of HKI-N to VDAC1 is disrupted by mutation of the membrane-buried Glu73. Moreover, we showed that mild acidification causes a selective and reversible dissociation of HKI-N from VDAC1–containing vesicles, consistent with our simulations and previous studies in intact cells. Our assay pinpoints a direct interaction of HKI-N with VDAC1, opening possibilities for systematic studies on the mechanisms that modulate HKI-VDAC complex assembly.

Our previous MD simulations indicated that membrane insertion of HKI occurs adjacent to the bilayer-facing Glu residue where a pair of polar residues mediates a marked thinning of the cytosolic leaflet [40]. This creates a gateway for the enzyme’s *N*-terminal α-helix to contact the negatively charged Glu residue and facilitate complex assembly. Therefore, it would be interesting to further explore the role of collective membrane properties such as lipid composition or membrane curvature, as well as channel density on HKI-N binding to VDAC1. These parameters could be readily varied in our assay, e.g. by using vesicles with increased diameter as compared to the very small unilamellar vesicles used in this study. Challenges due to compositional heterogeneity could be addressed by interrogation at single-vesicle and single-molecule level, which could be readily implemented by this assay format. Furthermore, the assay could be very useful for testing small therapeutic agents that neutralize pathogenic imbalances in this process [30, 32, 33].

Besides binding HKI, VDAC1 and notably VDAC2 function as dynamic translocation platforms for pro-apoptotic Bcl-2 proteins Bax and Bak [5, 11, 12], which regulate the permeability of the OMM for cytochrome *c* to initiate the apoptotic cascade. We previously reported that directing CERT-mediated ceramide transport to mitochondria induces Bax-dependent apoptosis [15, 16]. In subsequent studies, we identified VDAC2 as direct effector of ceramide-induced cell death and demonstrated that this function critically relies on the channel’s charged bilayer-facing Glu, which mediates direct contacts with the ceramide head group [17]. We here show that the same Glu residue of VDAC1 directly participates in HKI binding. The finding that ceramides share a common binding site with HKI on VDACs raises the possibility that ceramides exert their anti-neoplastic activities as modulators of VDAC-based platforms that control mitochondrial recruitment of pro-apoptotic Bcl-2 and anti-apoptotic HKI proteins. The vesicle-based VDAC-HKI interaction assay developed in this study opens exciting opportunities to experimentally challenge this model.

## METHODS

### Chemicals

L-α-phosphatidylcholine (eggPC) was purchased as chloroform solution from Avanti Polar Lipids (cat. no. 840051P) and 1,2-dihexadecanoyl-*sn*-glycero-3-phosphoethanolamine conjugated to Oregon Green 488 (DHPE-OG488) was from Invitrogen (cat. no. O12650). Maleimide-functionalized DY-547P1 and DY-647P1 were from Dyomics (cat. no. 647P1-03 and 547P1-03). Heptakis(2,6-di-O-methyl)-β-cyclodextrin was ordered from MP Biomedicals (cat. no. 157320). For surface functionalization, poly(ethylene glycol)diamine (DAPEG, cat. no. 753084), 3-glycidyloxypropyl trimethoxysilane (GOPTS, cat. no 440167) and 3-(maleinimido)-propionic acid N-hydroxysuccinimideester (NHS-maleimide, cat. no 63179) were ordered from Sigma Aldrich. HKI-N peptide (sequence: MIAAQLLAYYFTELKDDQCKKIDKY) was synthesized by Proteogenix (France). α-VDAC1 antibody was purchased from Cell Signaling (cat. no 4661, IB 1:1000). α-MBP antibody was purchased from Biolabs (cat. no E8030-s, IB 1:5000) and α-Rabbit-HRP was purchased from BioRad (cat. no 170-6515, IB 1:5000). Chemicals for buffers were purchased from AppliChem, Carl Roth or Sigma Aldrich. All aqueous solutions were prepared using ultrapure water, degassed and filtered through 0.2 μm hydrophilic membranes.

### DNA constructs

For expression of human VDAC1 and VDAC1^E73Q^ in *E. coli*, the corresponding cDNAs were PCR-amplified using Phusion high-fidelity DNA polymerase (Thermo Fischer Scientific) and inserted via *NdeI* and *XbaI* sites into bacterial expression vector pCold I (Takara Bio, USA). To produce single cysteine variants (C127S, A134C, C232S), the corresponding mutations were introduced using the QuickChange II site-directed mutagenesis method (Alignet, USA). A DNA fragment encoding αMBP-DARPin off7 with a *N*-terminal His6 tag and a *C*-terminal cysteine residue (αMBP-DARPin) [45] was subcloned via *NdeI* and *NotI* sites into the bacterial expression PET21a vector. The bacterial expression construct encoding MBP-TMD is described in [42]. Primers used for cloning and site directed mutagenesis of VDAC1 are listed in Supplementary Table 1.

### MD simulations

Coarse-grained [CG] MD simulations and simulation analysis were carried out according to Bieker *et al*. [40] Briefly, a refined solution NMR structure of VDAC1 (PDB: 6TIQ) and the first 18 residues of the *N-*terminal helix of a rat HKI structure (PDB: 1BG3), which is identical in sequence to the unresolved human HKI *N-*terminus, were coarse-grained using the martinize2 script [46] and then used for all CG MD simulations. VDAC1 was embedded into a 100 mol% POPC membrane, consisting of 770 lipids in total using the insane script [47] and approximately 150mM NaCl was added to the system, with an excess of Na^+^ to reach charge neutrality. For all simulations, the Martini 3 force field [48] with updated lipid models [49] was used, with the secondary structure of all proteins being restricted via an elastic-network approach [50]. Simulations were run with GROMACS versions 2021 [51]. All systems were minimized using a steepest descent algorithm. During equilibration, the HKI-N peptide was subjected to a flat-bottom harmonic restraint in z to ensure its close proximity to the bilayer and a harmonic restraint in x and y, to prevent any movement away from its initial placement at the box edge. Both restraints were lifted after equilibration. During the production run, the peptide was restrained by a repulsive harmonic potential within a vertical thickness of 1.5 nm at the z=0 position to prevent its desorption from the membrane, while simultaneously leaving its membrane-adsorbed state unaffected.

HKI-N-VDAC interactions were analyzed as contacts between the proteins’ particles within a 0.6 nm cutoff, grouped into contacts per residue. Residues were considered in contact if they have any particles in contact. These were plotted as either HKI-N Met1 contacts to any VDAC1 residue or as any HKI-N residue contacting VDAC1-E73 or VDAC1-Q73. Contact intensity is shown as the fraction of total simulation time for which the contact was established. Images of the HKI-N-VDAC interaction were produced using the Visual Molecular Dynamics (VMD) software [52] with either the all atom structures of VDAC1 (PDB: 6TIQ) and rat HKI (PDB: 1BG3) or as stills imaged directly from an MD simulation production run of VDAC1 with a deprotonated E73 and a protonated HKI-N *N-*terminus.

### Production of αMBP-DARPin

*E. coli* Bl21 (DE3) cells transformed with the αMBP-DARPin construct were grown in Lennox Broth medium (LB; tryptone 10 g/l, yeast extract 5 g/l, NaCl 5 g/l pH 7.0) containing 100 µg/m ampicillin to an OD_600_ of 0.4-0.6 prior addition of IPTG (0.5 mM) to induce protein expression. After growth was continued for 4 h at 37°C, cells were collected by centrifugation (5,350 x g, 15 min, 4°C) and lysed in Buffer A (20 mM Hepes pH 7.5) supplemented with protease inhibitor cocktail (PIC; 1 mg/mL apoprotein,1 mg/mL leupeptin, 1 mg/mL pepstatin, 5 mg/mL antipain, 157 mg/mL benzamidine) by microtip sonication with constant cooling in an ice-water bath. After centrifugation (25.000 x g, 25 min, 4°C), the lysate supernatant was loaded on a HiTrap Chelating HP column (Cytiva, USA) preloaded with Ni^2+^-ions and equilibrated in Buffer A. After washing the column with 5 column volumes of buffer A, a linear imidazole gradient (0-500 mM) was applied at a flow rate of 3 mL/min. Peak fractions were pooled and supplemented with 5 mM EDTA and 5 mM DTT while stirring on ice for 1 h. The affinity-purified protein was concentrated using an Amicon Ultra-4 unit (MWCO 10 kDa; Merck Millipore) and applied onto a HiPrep 16/60 200 HR Size exclusion chromatography column (Cytiva, USA) equilibrated in HBS (20 mM Hepes, 150 mM NaCl, pH 7.5) and fractionated at a flow rate of 1 mL/min. Peak fractions were pooled and the purified protein was stored at a concentration of 104 µM at −80°C.

### Production of MBP-TMD

*E. coli* Bl21 (DE3) cells transformed with the MBP-TMD construct were grown in LB medium containing 100 µg/m ampicillin to an OD_600_ of 0.4-0.6 prior addition of IPTG (0.5 mM) to induce protein expression. After growth was continued for 4 h at 37°C, cells were collected by centrifugation (5,350 x g, 15 min, 4°C) and lysed in 20 mL Lysis Buffer (150 mM NaCl, 20 mM Hepes, 1% Triton X-100, ∼2 mg DNase I, ∼5 mg Lysozyme, 1 x PIC, pH 7.5) by micro-tip sonication with constant cooling in an ice-water bath. After centrifugation (25.000 x g, 25 min, 4°C), the lysate supernatant was loaded on a HiTrap Chelating HP column (Cytiva, USA, catalog no. 17040801) precharged with Ni^2+^-ions and equilibrated in Loading Buffer (20 mM Hepes, 150 mM NaCl, 0.6 mM Triton X-100, pH 7.5). Loading was done in Loading Buffer containing 10 mM imidazole. After washing the column with 5 column volumes in the same buffer, a linear imidazole gradient (10-500 mM) in Loading Buffer was applied at a flow rate of 1 mL/min. Peak fractions were collected, pooled and loaded onto a Superdex 200 Increase 10/300 GL size exclusion column (Cytiva) pre-equilibrated in HBS buffer containing 0.6 mM TritonX-100. Peak fractions were pooled and the purified protein was stored at a concentration of 9.5 µM at −80°C.

### Production and cysteine-labelling of VDAC1

*E. coli* Bl21 (DE3) ΔOmp9 cells transformed with human VDAC1 (VDAC1, C127S, A134C, C232S) and VDAC1^E73Q^ (VDAC1, E73Q, C127S, A134C, C232S) expression constructs were grown in LB medium containing 100 µg/mL ampicillin. To induce VDAC1 protein expression, cells were grown to an OD_600_ of 0.4-0.6 at 37°C, cooled down for 30 min to 4°C, and then cultured in the presence of 1 mM of IPTG o/n at 15°C. Cells were collected by centrifugation (5,350 x g, 15 min, 4°C) and lysed in TEN Buffer (50mM Tris/HCl pH 8.0, 100mM NaCl) supplemented with protease inhibitor cocktail (PIC) through micro-tip sonication. Inclusion bodies were collected by centrifugation (30 min, 25.000 x g, 4°C), washed three times in TEN Buffer containing 2.5% Triton X-100, and then three times in TEN Buffer w/o Triton X-100. Inclusion bodies resuspended in TEN Buffer w/o Triton X-100 were diluted 1:10 by dropwise addition into 25 mM Na^+^PO4 pH 7.0, 100 mM NaCl, 6 M guanidine hydrochloride, 1 mM EDTA and 10mM DTT and then stirred o/n at 4°C. Next, the suspension was diluted 1:10 by dropwise addition to Refolding Buffer I (25 mM Na^+^PO_4_ pH 7.0, 100 mM NaCl, 1mM EDTA, 2.2% lauryldimethylamine oxide (LDAO)) and stirred o/n at 4°C. Finally, the suspension was diluted 1:10 by dropwise addition to Refolding Buffer II (25 mM Na^+^PO_4_ pH 7.0, 10 mM NaCl, 1mM EDTA, 0.1% LDAO), stirred overnight at 4°C, passed through a 0.2 µm filter, degassed and loaded on a Fractogel EMD-SE Hicap cation exchange column (Merck Millipore, USA). After washing the column with 5 column volumes of Refolding Buffer II, a linear salt gradient (10-1000 mM NaCl) was applied at a flow rate of 1 mL/min. Peak fractions were collected, pooled and exchanged against SEC Buffer (10mM Tris/HCl, pH 7.0, 100mM NaCl, and 0.05% LDAO) using a PD column (Cytiva). For cysteine labelling, 35 µM of purified protein were mixed with 105 µM DY-547P1 for 45 min in the dark at RT. After addition of 315 µM L-cysteine, the mixture was incubated for 15 min at RT and centrifuged (10,000 x g, 10 min, 4°C). The supernatant was loaded onto a Superdex 200 Increase 10/300 GL size exclusion column (Cytiva) and eluted in SEC Buffer. Protein purity and efficiency of cysteine-labeling was determined by SDS-PAGE, colloid Coomassie Blue staining and in-gel fluorescence (IGF) analysis using a Typhoon FLA 9500 Biomolecular Imager (GE Healthcare Life Sciences, USA) with a 555 nm excitation laser, LPB filter, 50 µm pixel size, and PMT voltage set to 290 V. Purified proteins were snap-frozen and stored at −80°C at a concentration of 140 µM (VDAC1), 106 µM (VDAC1^E73Q^), 32 µM (DY-547P1-VDAC1, 71% labeled) and 13 µM (DY-547P1-VDAC1^E73Q^, 80% labeled).

### Surface modification and functionalization with αMBP-DARPin

Standard microscopy glass coverslips (24 mm diameter, 1.5 mm thickness) were covalently modified with a dense PEG polymer brush and functionalized with maleimide for immobilizing αMBP-DARPin using previously established protocols [53, 54]. First, the cover slips were cleaned by sonication in acetone for 15 min followed by sonication in double-deionized water (DDW) for 15 min, dried under a stream of N_2_, and then plasma cleaned in ambient air for 10 min. The cleaned surfaces were then assembled into sandwiches with 10 µL GOPTS and incubated for 45 min at 75 °C for silanization.

Next, the surfaces were washed in acetone and dried under a stream of N_2_. Half of the silanized coverslips were covered with a spatula tip of DAPEG (approx. 10 mg) and again assembled into sandwiches with the other half of the coverslips. The sandwiched pairs were incubated for 20 min at 75°C to melt the DAPEG. The upper cover slip was moved to ensure the entire glass surface was evenly coated with DAPEG and remove air bubbles. For proper PEGylation, the sandwiched pairs were incubated for 4-20 h at 75°C followed by washing in dH_2_O and 5 min of sonication in dH_2_O. In the last step, the cover slides were dried by a stream of N_2_. GOPTS-PEG functionalized coverslips were either used directly for NHS-maleimide functionalization or stored at 4 °C until use. GOPTS-PEG surfaces were further functionalized using NHS-maleimide. To this end, maleimide was dissolved in DMF at a final concentration of 0.05 µg/µL and 9 µL were applied to a coverslip, which was then “sandwiched” with a second one. After incubation for 30 min at RT, the coverslips were washed in chloroform, dried under a stream of N_2_, and incubated with 10 µM αMBP-DARPin in HBS (150 mM NaCl, 20 mM Hepes) pH 6.5 for 1 h at RT. The coverslips were washed three times in HBS pH 7.5 before use.

### Co-reconstitution of MBP-TMD and VDAC1 and application to modified surface

Membrane protein reconstitution into very small unilamellar vesicles (VSUV) was achieved by detergent extraction using methyl-β-cyclodextrin as previously described [42, 55]. To this end, 0.25 mM of lipid (egg PC:DHPE-OG488, 99.5:0.5, mol%) was mixed with MBP-TMD and VDAC1 or VDAC1^E73Q^ in HBS pH 7.5 containing 22 mM TX-100 at a molar protein:lipid ratio of 1:500, 1:50 and 1:100, respectively, in a total reaction volume of 500 µL. After incubation for 1 h at RT, methyl-β-cyclodextrin was added to the reconstitution mixture in a twofold excess (molar ratio) over the detergent and incubated for 5 min at RT. The reaction mixture was filled up to a final volume of 500 µL with HBS pH 7.5 and applied to the functionalized cover slips. After 30 min of incubation (at RT), the cover slips were washed in HBS pH 7.5 by pipetting up and down several times all over the surface to remove unbound vesicles. After two additional washing steps in HBS pH 7.5, the functionalized coverslips with bound vesicles were ready for TIRF microscopy and fluorescence recovery after photobleaching (FRAP) measurements.

### FRAP measurements and data analysis

HK-N peptide was labelled at a concentration of 10 µM with 30 µM DY-647P1 in HBS buffer for 45 min in the dark at RT, followed by incubation with 90 µM L-cysteine for 15 min in the dark at RT to block the unreacted dye. DY-647P1-labeled HK-N peptide (200 nM or 400 nM) was applied in HBS pH 7.5 to the surface of functionalized coverslips with bound vesicles and incubated for 5 min at RT prior to fluorescence microscopy and FRAP measurements. Image acquisition was performed in total internal reflection fluorescence (TIRF) illumination using an inverted Olympus IX-83 microscope. The setup included an sCMOS camera (ORCA-Flash 4.0, Hamamatsu), a 4-line TIRF condenser (Olympus) fiber-connected to a 488 nm laser diode (200 mW, LuxX+), a 561 nm fiber laser (50 mW, MBP communications), and a 642 nm fiber laser (50 mW, MBP communications), as well as a 405 nm laser diode for FRAP (60 mW, LuxX+). A UPLAPO 100 x HR objective with a NA of 1.5 and oil immersion was used for TIRF illumination and image capturing. Precise positioning was ensured by a motorized ultrasonic xy-stage (IX3-SSU, Olympus). Shifts in the z-plane were prevented by a hardware autofocus (IX3-ZDC2, 830 nm version, Olympus). For FRAP measurements, the cellFRAP (Olympus) unit was used with the following bleaching set up: for bleaching of lipid-associated (OG488) or VDAC1-associated (DY-547P1) fluorescence, the 405 nm laser was set to 50% laser power and the sample was bleached for 200 ms followed by 10 cycles of image acquisition with a 30 s interval between each image acquisition. For FRAP measurements in the DY-647P1 channel (642 nm), the 405 nm laser was set to 50% laser power and the sample was bleached for 200 ms followed by 150 cycles of image acquisition with a 0.5 s interval between each image acquisition. To increase the fluorescence intensity, a 2×2 binning was applied. Data analysis and processing was done with FIJI (ImageJ NIH, Bethesda, MD). For FRAP curves, analysis was done with a bleaching correction using the following equation:

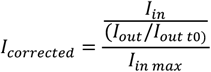

For quantifying *I*_*in*_, in a circular ROI was drawn into the region of bleaching. Intensity out was determined by a circular ROI outside of the bleaching region to measure the overall fluorescence decrease. For analysis, the FIJI plugin Time Series Analyzer v3 was used. For intensity plots, the offset of the 2×2 binning (400 counts) was subtracted. To determine the lifetime τ of the interaction, a mono exponential decay was fitted from 1-25 s (for pH 6.5 0.5-10s) of the curves using the equation: *I* = *I*_0_ + *A*_1_*e*^−*t*/*τ*^ using Origin (OriginLab, USA)

### Quantitative assessment of VDAC1 vesicles captured on modified surface

Following FRAP experiments, vesicles captured on functionalized glass substrates were extracted by application of 200 µl of 1 x Laemmli Buffer (50% Glycerol, 10% SDS, 10% β-mercaptoethanol, 0.025 % Bromophenolblue, 0.3 M Tris/HCl pH 6.8) prewarmed at 60°C. After 2 min of incubation, the coverslip was rinsed in the same solution by rigorous pipetting. This procedure was repeated three times with the same solution. Next, vesicle extracts were subjected to immunoblot analysis using antibodies against VDAC1 and MBP.

## ACKNOWLEDGEMENTS

This work was supported by the Deutsche Forschungsgemeinschaft (378148610, 448344643 and 467522186/SFB1557–Z1 to J. C. M. H.; 467522186/SFB1557–P13 to J.P.) and the FCT – Fundação para a Ciência e a Tecnologia I.P. (through MOSTMICRO-ITQB R&D Unit with projects UIDB/04612/2020 and UIDP/04612/2020, and LS4FUTURE Associated Laboratory with projects LA/P/0087/2020 and CEECIND/04124/2017/CP1428/CT0008 to M. N. M.).

## Author Contributions

J. P. and J. C. M. H. designed the research; M. W. performed experiments with critical input from I. W. and B. F.; M. T. carried out the CG-MD simulations with critical input from M. N. M.; J. P., I. W. and J. C. M. H. provided expertise for experiments and helped interpret the data; M. N. M. provided expertise for CG-MD simulations and helped interpret the data; M. W. and J. C. M. H. wrote the manuscript; all authors discussed results and commented on the manuscript.

## Competing Interests

The authors declare no competing interests.

## Data Availability

All data generated or analyzed in this study are included in the manuscript and supporting files. Source data with sample sizes, number of technical and/or biological replicates, means, standard deviations, and calculated *p* values (where applicable) are provided in the Supplementary Data file. Uncropped scans of immunoblots are provided in Supplementary Information.

## Conflict of interests

The authors declare that they have no conflicts of interest with the contents of this article.

## Data availability

All data generated in this study are included in the manuscript and Supplementary Information file.

## Notes

### Competing Interest Statement

The authors have declared no competing interest.

