## Supplemental Information for "Key determinants of VDAC-hexokinase I complex assembly revealed by a minimal vesicle-based interaction assay"

**Affiliations:**
[1]Division of Molecular Cell Biology, Department of Biology/Chemistry, Osnabrück University, 49076 Osnabrück, Germany.
[2]Division of Biophysics, Department of Biology/Chemistry, Osnabrück University, 49076 Osnabrück, Germany.
[3]Center for Cellular Nanoanalytics, Osnabrück University, 49076 Osnabrück, Germany.
[4]Instituto de Tecnologia Química e Biológica António Xavier, Universidade Nova de Lisboa, Av. da República, 2780-157 Oeiras, Portugal.

**This PDF file includes:**

Supplementary Figs. 1 to 8

Supplementary Table 1

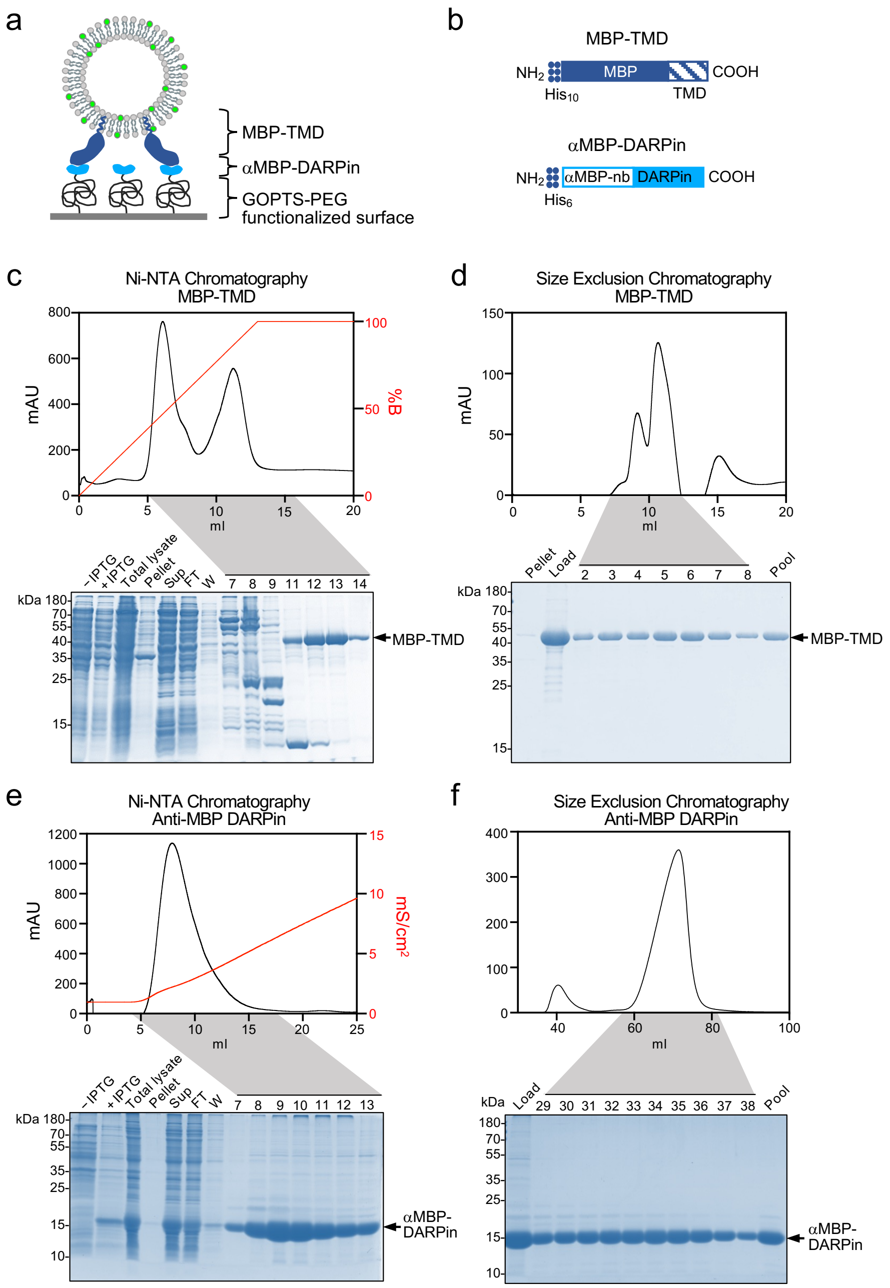


**Supplementary Figure 1 | Purification of MBP-TMD and αMBP-DARPin.**

(**a**) Cartoon of vesicle containing maltose binding protein fused to a transmembrane domain (MBP-TMD) captured on a glass coverslip decorated with αMBP-DARPin via crosslinking to GOPTS-PEG-NHS maleimide.

(**b**) Schematic outline of the MBP-TMD and αMBP-DARPin proteins used for capturing vesicles to glass surface as in (a).

(**c**) Ni-NTA chromatography of MBP-TMD upon its IPTG-induced expression in *E. coli*. After binding to the Ni-NTA matrix, MBP-TMD was released using a linear imidazole gradient. Expression, release from bacterial lysates, and Ni-NTA chromatography of MBP-TMD was monitored by SDS-PAGE and CB staining. Sup, supernatant; FT, flow through.

(**d**) MBP-TMD affinity-purified by Ni-NTA chromatography was loaded onto a size exclusion column. Peak fractions were analyzed by SDS-PAGE and CB staining.

(**e**) Ni-NTA chromatography of αMBP-DARPin upon its IPTG-induced expression in *E. coli*. After binding to the Ni-NTA matrix, αMBP-DARPin was released using a linear imidazole gradient. Expression, release from bacterial lysates, and Ni-NTA chromatography of αMBP-DARPin was monitored by SDS-PAGE and CB staining.

(**f**) αMBP-DARPin affinity purified by Ni-NTA chromatography was loaded onto a size exclusion column. Peak fractions were analyzed by SDS-PAGE and CB staining.

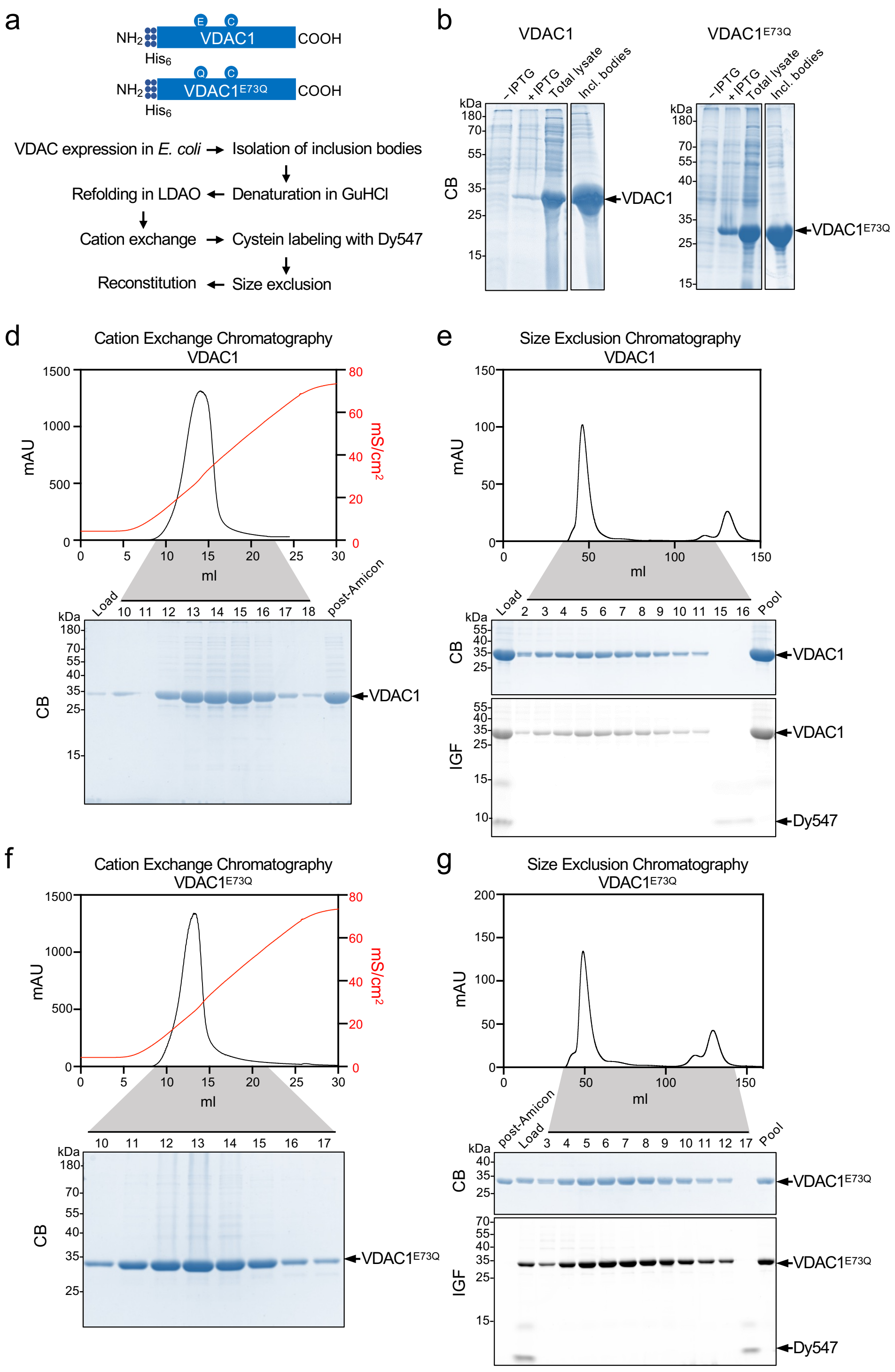


**Supplementary Figure 2 | Purification and cysteine labelling of VDAC1 and VDAC1$^{E73Q}$**

(**a**) Schematic outline of the production, purification and cysteine-labeling of recombinant human VDAC1 using an IPTG-inducible bacterial expression construct encoding single cysteine VDAC1 (C127S, A134C, C232S) or its HKI binding-deficient variant VDAC1$^{E73Q}$ (E73Q, C127S, A134C, C232S) tagged with a *N*-terminal His-tag.

(**b**) Inclusion bodies isolated from *E. coli* following IPTG-induced expression of VDAC1 or VDAC1$^{E73Q}$ were analyzed by SDS-PAGE and CB staining.

(**c**) VDAC1 solubilized from inclusion bodies was refolded, loaded onto a cation exchange column and released using a linear salt gradient. Cation exchange chromatography of VDAC1 was monitored by SDS-PAGE and CB staining. Peak fractions were pooled and concentrated using an Amicon filter (post Amicon).

(**d**) VDAC1 purified by cation exchange chromatography was cysteine-labeled with DY-547P1 and loaded onto a size exclusion column. Peak fractions were analyzed by SDS-PAGE, CB staining and in-gel-fluorescence (IGF) analysis.

(**e**) Cation exchange chromatography of VDAC1$^{E73Q}$ performed as in (c).

(**f**) Size exclusion chromatography of Dy-547P1-labelled VDAC1$^{E73Q}$ performed as in (d).

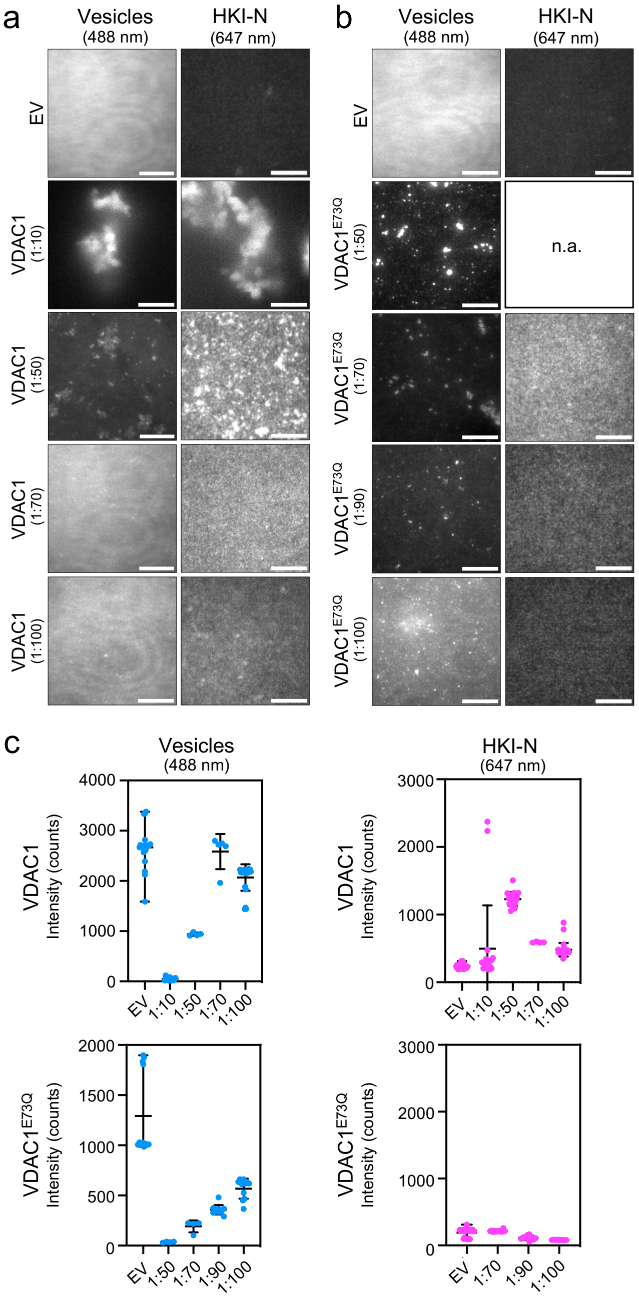


**Supplementary Figure 3 | Protein-lipid ratio optimization for reconstitution of MBP-TMD and VDAC1 in proteoliposomes**.

(**a, b**) Vesicles containing MBP-TMD in combination with VDAC1 or VDAC1$^{E73Q}$ were prepared using molar protein:lipid reconstitution ratios of 1:500 for MBP-TMD and the indicated ratios for VDAC1 and VDAC1$^{E73Q}$ in eggPC:DHPE-OG488 (99.5:0.5). Vesicles were captured on αMBP-DARPin-decorated coverslips, incubated with 200nM HK-N$^{647}$ and analyzed for OG488 and HK-N$^{647}$ fluorescence using TIRF microscopy.

(**c**) Intensity plots of OG488 (*blue*) and HK-N$^{647}$ (*magenta*) fluorescence at the surface of coverslips as in (a, b). Data shown are mean values ± SD.

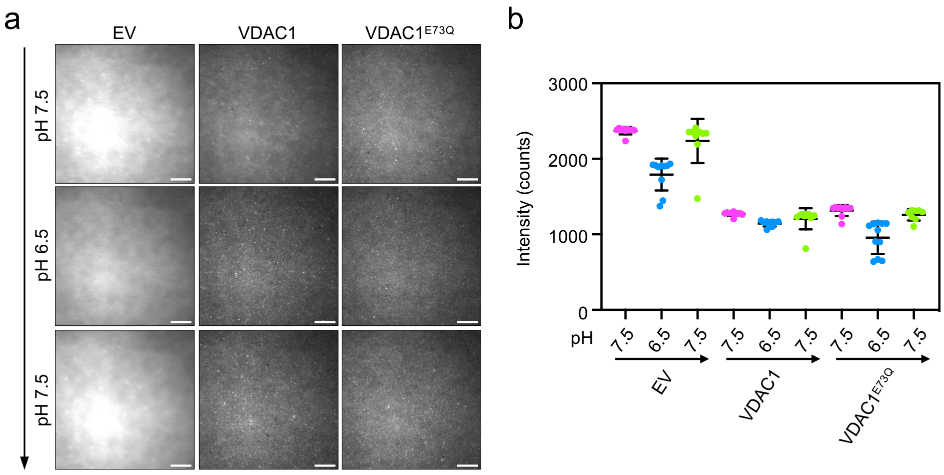


**Supplementary Figure 4 | Mild acidification does not affect binding of VDAC-containing vesicles to αMBP-DARPin-decorated coverslips**.

(**a**) Vesicles prepared by (co)-reconstitution of MBP-TMD, VDAC1 and/or VDAC1$^{E73Q}$ at molar protein:lipid reconstitution ratios of 1:500 (MBP-TMD), 1:50 (VDAC1) and 1:100 (VDAC1$^{E73Q}$) in eggPC:DHPE-OG488 (99.5:0.5) were captured on αMBP-DARPin-decorated coverslips in pH 7.5 buffer. After 5 min, OG488 fluorescence levels were measured by TIRF microscopy. Incubation then continued in pH 6.5 buffer for 5 min before measuring OG488 fluorescence levels again. This procedure was then repeated once more in pH 7.5 buffer.

(**b**) Intensity plots of OG488 fluorescence at the surface of coverslips treated as in (a). Data shown are representative of three independent experiments and plotted as mean values ± SD of 5-10 different positions per condition.

**Supplementary Table 1. Primers used for cloning and site-directed mutagenesis.**

| Primer name | Primer sequence (5’-3’) |
|---|---|
| pColdI-hVDAC1 F | ATCATATCGAAGGTAGGCACGCTGTGCCACCCAC |
| pColdI-hVDAC1 R | GCTTTTAAGCAGAGATTACCTATTTATGCTTGAAATTCCAGTCCTAGACCAAG |
| hVDAC1-E73Q F | ATGGACTGAGTACGGCCTGACGTTTACACAGAAATGGAATACCGAC |
| hVDAC1-E73Q R | GTCGGTATTCCATTTCTGTGTAAACGTCAGGCC |
| hVDAC1 A134C F | TTTCGACATTTGCGGGCCTTCCATCCGGGG |
| hVDAC1 A134C R | TCCATGTCGCTGCCCAGG |
